# Myopia alters the structural organization of the retinal astrocyte template, associated vasculature and ganglion layer thickness

**DOI:** 10.1101/2022.02.22.481546

**Authors:** Carol Lin, Abduqodir Toychiev, Nefeli Slavi, Reynolds Ablordeppey, Miduturu Srinivas, Alexandra Benavente-Perez

## Abstract

**Purpose:** To describe the effect of myopic eye growth on the structure and distribution of astrocytes, vasculature and ganglion cell thickness, critical for inner retinal tissue homeostasis and survival.

**Methods:** Astrocyte and capillary distribution, retinal nerve fiber (RNFL) and ganglion cell layer (GCL) thicknesses were assessed using immunochemistry and spectral domain optical coherence tomography on eleven retinas of juvenile common marmosets (*Callithrix Jacchus*), six of which were induced with lens-induced myopia (refraction, Rx: −7.01±1.8D). Five untreated age-matched juvenile marmoset retinas were used as controls (Rx: −0.74±0.4D).

**Results:** As control marmoset eyes grew normally, there was an age-related increase in astrocyte numbers associated with RNFL thickening. Marmosets with induced myopia did not show this trend and, on the contrary, had reduced astrocyte numbers, increased positive GFAP immunopositive staining, thinner RNFL, lower peripheral capillary branching, and increased numbers of string vessels.

**Conclusion:** The myopic changes in retinal astrocytes, vasculature, and ganglion cell layer thickness suggest a reorganization of the astrocyte and vascular templates during myopia development and progression. Whether these adaptations are beneficial or harmful to the retina remains to be investigated.

**Summary Statement:** This article provides new information on how progressive myopia affects key elements of the retinal neurovascular unit.

## INTRODUCTION

Myopia is a significant risk factor for glaucoma, maculopathy, and choroidal neovascularization among others(Saw et al., 2005, Curtin, 1985). All myopes, regardless of degree, are at increased risk of visual impairment(Curtin, 1985, Saw et al., 2005, Xu et al., 2007, Foster and Jiang, 2014). Despite the predicted global increase in myopia prevalence and its potential public health crisis in vision care(Curtin, 1985, Holden et al., 2016), the mechanisms that lead to myopic degeneration and associated conditions remain unknown. There is a lack of early diagnostic markers for myopic pathology and no means to prevent disease progression(Holden et al., 2016, Yokoi and Ohno-Matsui, 2018).

The development and maintenance of a healthy retina relies on the neurovascular interplay between neuronal, vascular and glial cells(Hawkins and Davis, 2005), which provide structural and nutritional support(Hawkins and Davis, 2005, Provis et al., 1997), regulate metabolism(Hawkins and Davis, 2005, Vecino et al., 2016, Sapieha, 2012), neuronal debris phagocytosis(Hawkins and Davis, 2005, Vecino et al., 2016, Sapieha, 2012), and ion and neurotransmitter homeostasis(Sapieha, 2012). The neurovascular unit exerts a biphasic influence of degenerative and subsequent reactionary regeneration in systemic pathology, that can be harmful at the acute phase and beneficial at the chronic phase(Maki et al., 2013).During normal development and aberrant disease process, blood vessels, retinal astrocytes and ganglion cells are in a reciprocal feedback loop(Ramirez et al., 1996). For example, astrocyte numbers and distribution are determined by retinal capillary density and amount of RNFL damage due to pathology(Dorrell et al., 2002, West et al., 2005). Astrocyte reactivity and degeneration precedes ganglion cell degeneration and pathological neovascularization, respectively(Dorrell et al., 2010, Sun and Jakobs, 2012).

Myopic eyes experience blur because they are larger in size, which appear to result in a compromised vascular support to the inner retina. Lower central retinal artery blood velocities(Benavente-Perez et al., 2010, Leng et al., 2018), narrowing of the retinal vessel diameter(Leng et al., 2018), decreased capillary density(Holden et al., 2016), and larger foveal avascular zones(Golebiewska et al., 2019) have been described in human myopic eyes with no degeneration. Due to the interrelationship between the neurovascular elements, these vascular changes might in turn compromise vascular and neuronal function, premeditate astrocytic reorganization and be part of the etiological cascade of events that increases the risk to develop posterior pole complications(Al-Sheikh et al., 2017, Zhu et al., 2020). However, the longitudinal effect of myopia on capillaries, astrocytes and ganglion cells, and how they relate to each other, remains unknown.

Research with non-human primates (NHP) is a critically important link between animal studies and human treatments(Kishi et al., 2014, Okano et al., 2012, Mansfield, 2003). The common marmoset (*Callithrix jacchus*) has been established as an excellent NHP model for vision and neuroscience research because of its small size, fast development, ease in breeding and handling, and high optical quality eye with diurnal foveated retina(Troilo and Judge, 1993, Nickla et al., 2002, Benavente-Perez et al., 2014). Common marmosets have successfully been induced with moderate and high myopia using negative contact lenses, following our well-established lens-induced myopia paradigm(Benavente-Perez et al., 2012, Benavente-Perez et al., 2014, Benavente-Perez et al., 2019, Nickla et al., 2002, Troilo and Judge, 1993). The strengths of our marmoset myopia model are the nonexistence of another NHP model of myopia that uses contact lenses to induce myopia, the controlled experimental conditions, and the ocular anatomy and physiology directly comparable to human eyes. In this study, the structure and distribution of three key elements of the retinal neurovascular unit – astrocytes, co-localized superficial capillaries, and RNFL/GCL thickness - were studied in a NHP model of myopia to assess the neurovascular changes that eyes experience during myopia development and progression and how they relate to each other, with the ultimate goal to understand the effect of progressing myopia on the ocular tissues.

## RESULTS

### Characterization of superficial capillaries and associated astrocytes in control marmoset retinas

Six myopic marmoset eyes were studied (age 200.3±14.2 days). Five untreated age-matched juvenile marmosets were used as controls (age 232.2±32.9 days). In control eyes, blood vessels were similar in width and shape within the fovea and parafovea (Figure 1B). The marmoset retina exhibited four vascular plexi in the foveal and parafoveal retina (Figure 2A, area 1 and 2): the radial peripapillary capillary plexus (RPC), superficial capillary plexus (SUP), intermediate capillary plexus (INT), and deep capillary plexus. The peripapillary retina had three vascular plexi (Figure 2A, area 3), while the peripheral retina had two vascular plexi (Figure 2A, area 4). The four vascular plexi had varying vessel diameters, morphology, and distribution (Figure 2B), with vasculature becoming denser and more regular in shape with deeper plexi.

**Figure 1:**
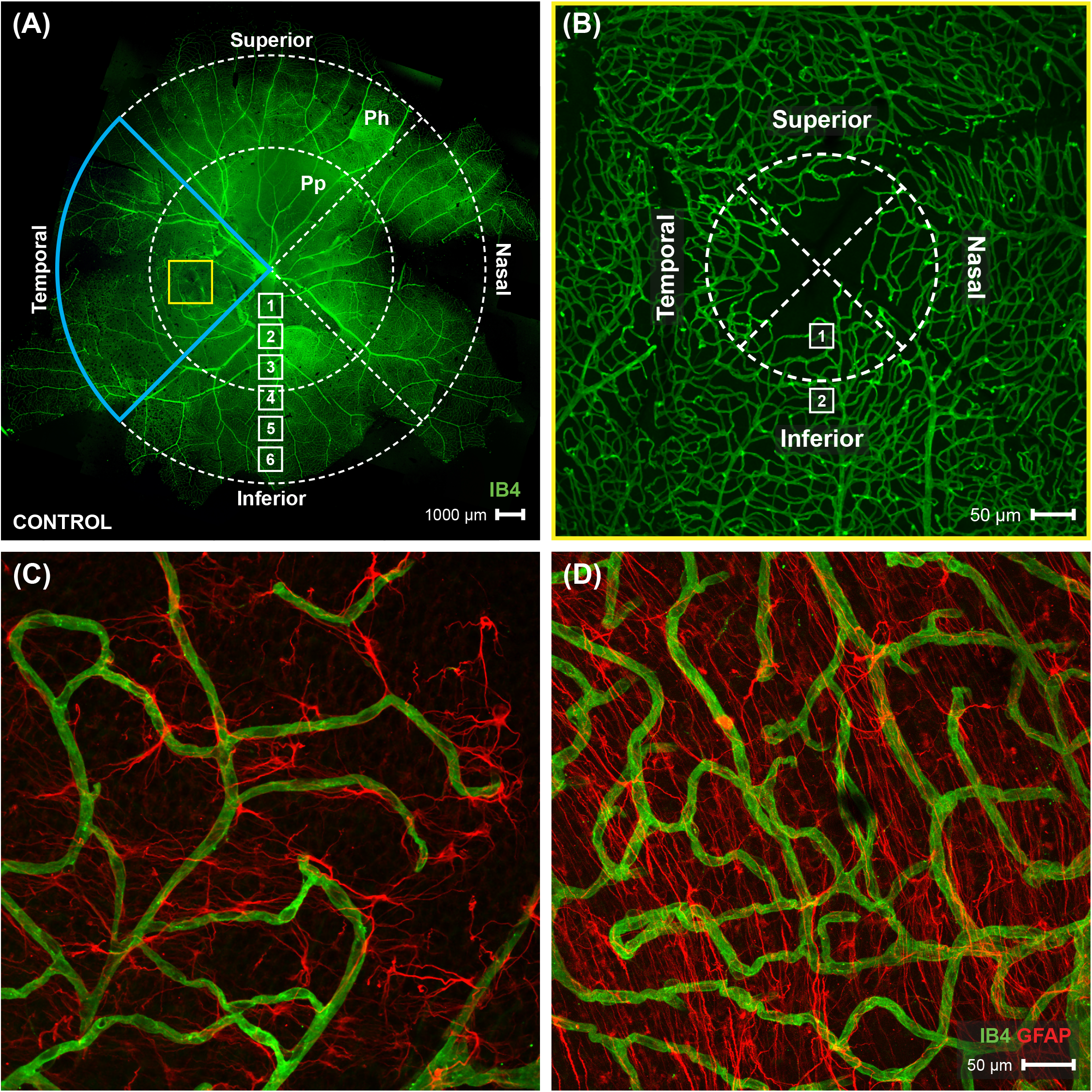
A map of the vasculature of the whole mount marmoset retina, and images of foveal and parafoveal region. **(A)** Shows a complete map of the retinal vasculature (green) obtained from control marmoset (ID: X15, age: 381 days, refractive error: −1.12D). Retinal vasculature was visualized with conjugated IB4-488. Images were acquired at 4x magnification and stitched in Photoshop. Location of peripapillary region (Pp) and peripheral region (Ph) described in this study is shown. Yellow box represents the foveal region, seen in panel 1B. White boxes represent focal areas away from the optic disc to periphery. Boxes labeled “A” represent location where peripapillary location images were taken, while boxes labeled “B” represent location where peripheral location images were taken. Superior, Inferior, nasal, and temporal quadrants of the retina are shown. **(B)** A representative image of the vasculature (green) in the foveal region of the control marmoset (ID: H16, age: 205 days, refractive error: −1.15D), taken at 10x magnification. Box labeled “1” represents the foveal region, while box labeled “2” represents the parafoveal location. **(C and D)** Shows anatomy of the astrocytes (red) and vasculature (green) in the fovea (1B, box1) and parafovea (1B, box2) region. Images were acquired at 40x magnification and astrocytes were visualized with GFAP marker.

**Figure 2:**
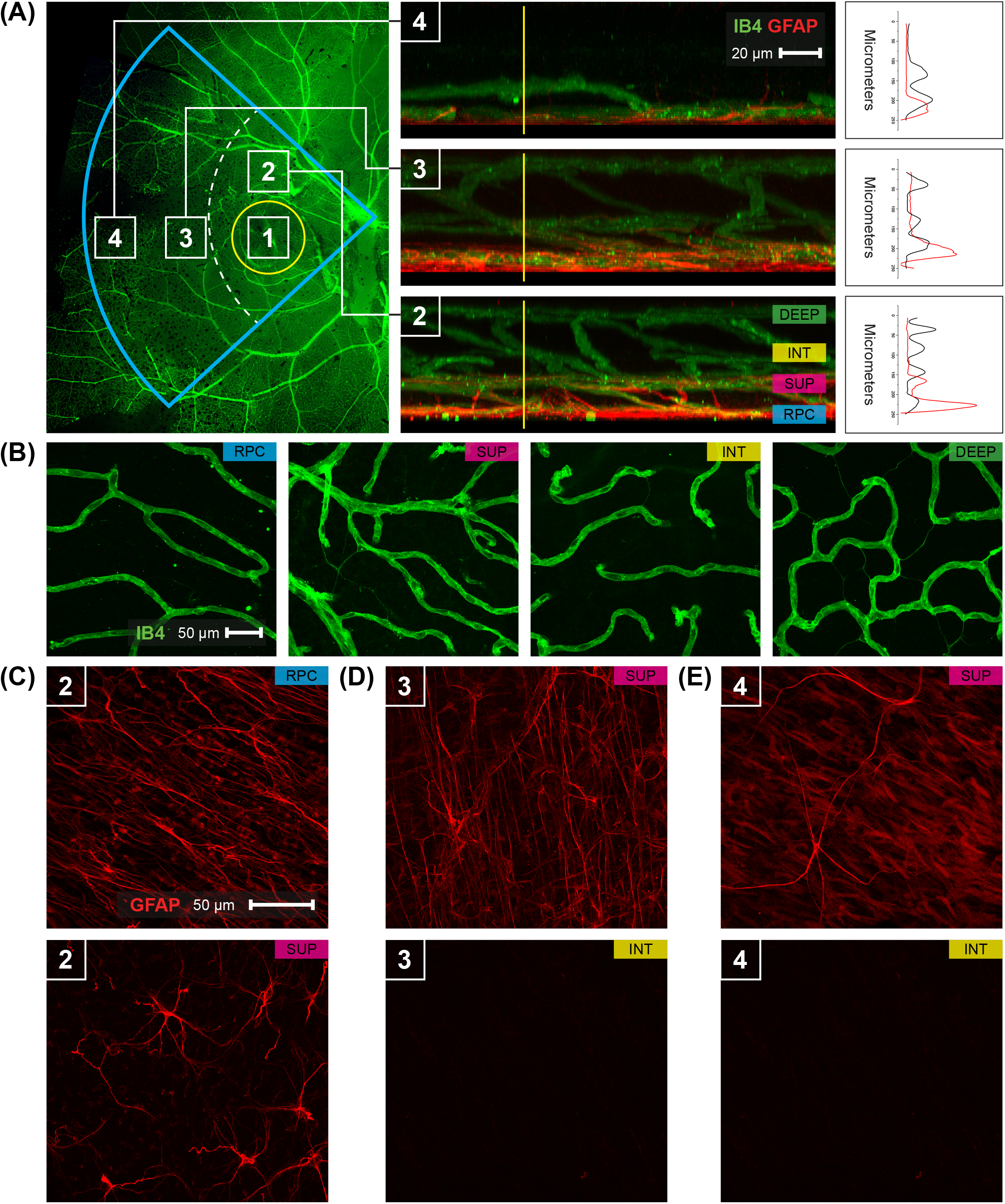
Retinal vasculature and astrocytes distribution in the temporal part of the control marmoset. **(A)** An image of the temporal part of the retina (left) is visualized with isolectin (green) and highlighted in a blue insert. Numbers in the white boxes represent areas (1: foveal, 2: parafoveal, 3: peripapillary, 4: periphery) that are used for 3D reconstructions. Reconstructed images (middle) from areas 2, 3, and 4 show the distribution of the inner vasculature (green) and astrocytes (red). Scale, 20 μm. A vertical line (yellow) is an intensity profile (right) used to show colocalization vasculature (black) with astrocytes (red) in GCL. **(B)** Representative images of the retinal vasculature (green) acquired from the parafoveal area show all vascular layers. Scale, 50 μm. **(C)-(E)** An image of the retinal astrocytes (red) in GCL that shows the morphology and distribution in areas 2, 3, and 4. Scale, 50 μm. In area 2 (parafovea) astrocytes differ in morphology and are distributed in two vascular layers RPC and SUP layers. In other areas (3, 4) areas astrocytes are found only in the superficial layer.

There were two layers of astrocytes in the foveal and parafoveal marmoset retina (Figure 2C), corresponding to the RPC and superficial vascular plexi. The astrocytes of the RPC were more numerous, compact, and elongated than those in the superficial layer which are less in number, less compact, and more stellate (Figure 2C). In the peripapillary and peripheral marmoset retina, there was only one layer of astrocytes (Figure 2D and 2E). There were decreasing amounts of astrocytes that become more stellate away from the optic nerve head and towards the periphery. The vasculature in focal areas closer to the optic nerve head contained vessel diameters of varying widths (Figure 3A, panel 1-3). The further away from the optic nerve head, the smaller and more uniform the diameters of the retinal blood vessels (Figure 3A, panels 4-6). String vessels were identified in the superficial capillary plexus.

**Figure 3:**
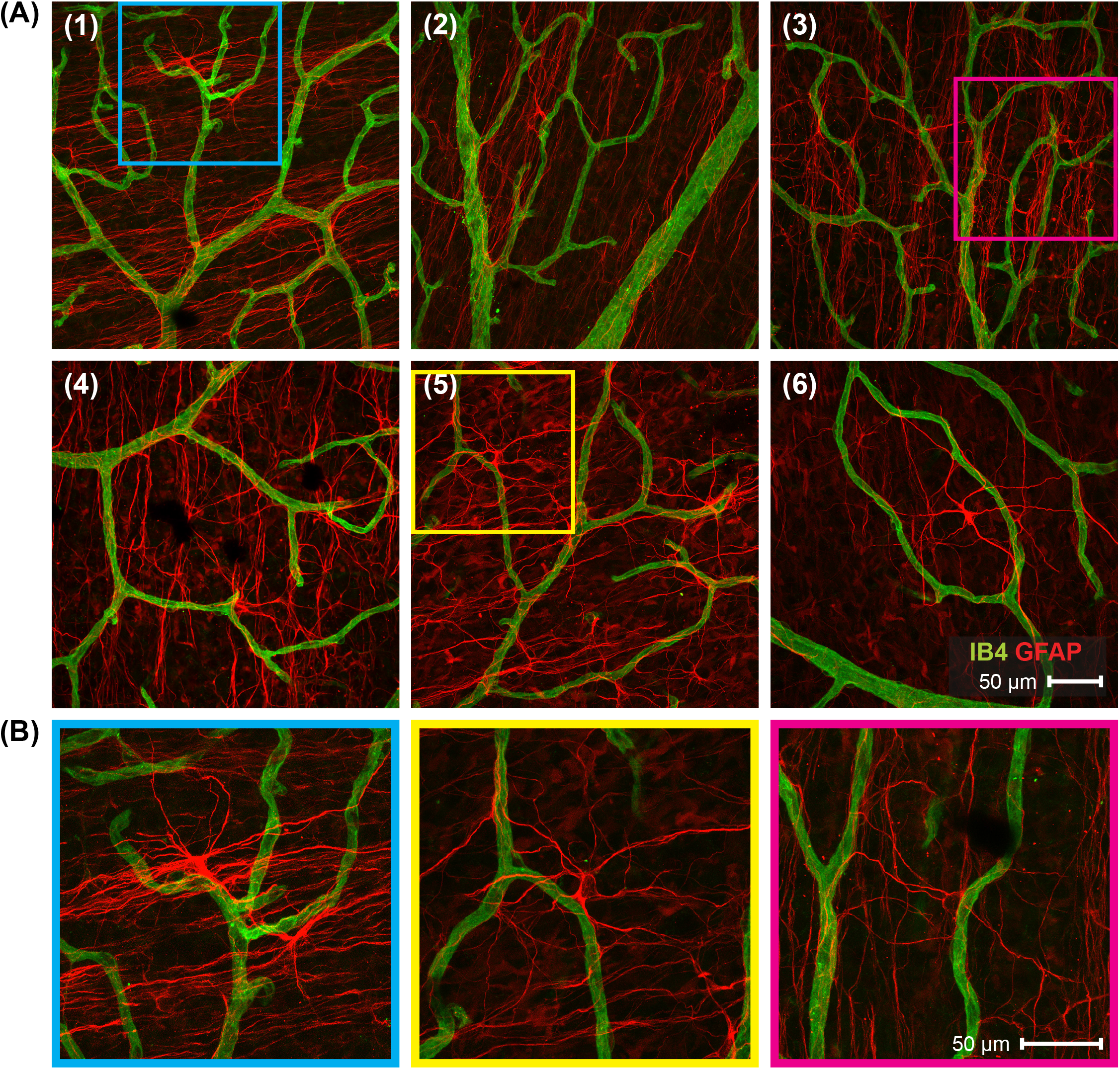
Distribution of the superficial astrocytes and vasculature in the marmoset retina, through visualization of focal areas from near the optic nerve head to the far periphery. **(A)** Representative images of the marmoset retina taken from superficial layer that shows distribution of the vasculature (green) and astrocytes (red) in focal areas as seen in figure 1A (focal area from the optic nerve head is denoted in the top left of each panel). Images were acquired from the control marmoset (ID tag: X15, age: 381 days old, Rx: −1.12 D) at 40x magnification. **(B)** Morphology of the astrocytes from specific areas are shown in colored boxes (blue in panel 1; pink in panel 3; yellow in panel 5). Magnified images demonstrate that astrocytes appear more elongated and denser in the areas 1 to 4 and at areas (5 and 6) they become stellate and exhibited less density. Organization and morphology of astrocytes at Superior, Inferior and Nasal sides were similar in control marmosets. Each panel represents a typical finding from a sample of five control marmosets.

Elongated astrocytes had smaller sized bodies than stellate astrocytes and were polygonal in shape. GFAP immunopositive staining per astrocyte was increased in the fovea, parafovea, and the peripapillary regions of the retina. The astrocytes in these regions were more dense, elongated and linear as they exited the optic nerve head (Figure 3A, panels 1-3), and became scarcer and more stellate as they approached the periphery (Figure 3A, panels 4-6). Representative astrocytes from control eyes can be seen in Figure 3B within panels 1, 3, and 5. These findings in control marmosets describe physiological variations in vessel and astrocyte anatomy expected to be found in the optic nerve head, fovea, and periphery of untreated marmoset retinas.

### Superficial capillaries branch less in the retinal peripapillary and periphery of myopic marmosets

Representative images of the superficial vasculature are shown in Figure 4A. Myopic eyes had significantly less branching in the periphery and peripapillary region, and increased branching in the foveal retina, compared to controls (all p<0.01; Figure 4B). There were also a greater number of string vessels in the myopic parafoveal superficial capillary plexus than in controls (<0.01)

**Figure 4:**
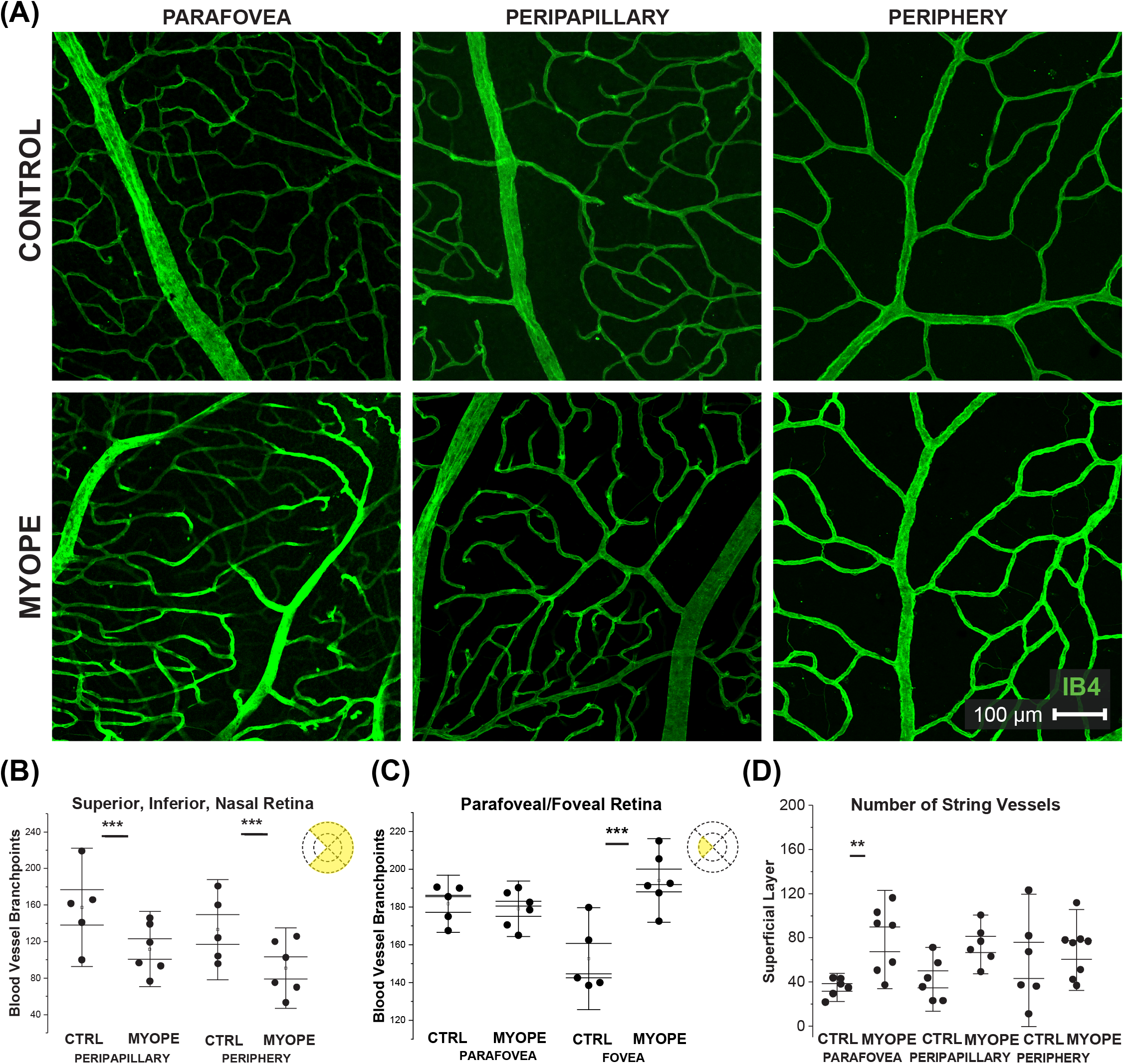
Vascular alterations of the myopic marmoset retina, shown in focal areas 2, 4, and 6 from the optic nerve head. **(A)** Representative images of superficial capillary structure in the parafoveal, peripapillary, and peripheral regions of a control marmoset (ID tag: X15, age: 381 days, Rx: −1.12D) and myopic marmoset (ID tag: U17, age: 179 days, Rx: −3.09D), taken at 20x magnification. Vasculature is labeled with IB4 (green). **(B)** Analysis of the capillary branch points in the peripapillary region and peripheral superficial vascular layer of the superior, inferior, and nasal retina. Data shown as a I-graph box plot for control (n=5) and myopic (n=6) marmoset retinas where the inner box lines represents standard error (SE) and outer lines/whiskers represents standard deviation (SD). The number of branch points were significantly different in the peripapillary and peripheral regions. Peripapillary P<0.01, Peripheral P<0.01 **(C)** Analysis of the capillary branch points in the fovea and parafovea region (control n=5, myope n=6). A significant increase in branching was seen in the myopic eye in the foveal regions, with no change noted in the parafoveal myopic retina. Fovea P<0.01, Parafovea P=0.16. **(D)** Analysis of the number of string vessels in the superficial capillary plexus (SCP). A significant increase in the number of string vessels in the myopic parafoveal SCP was noted (P<0.01). No significance was found in the other retinal areas (Peripapillary P=0.11, Periphery P=0.62).

### Myopic eyes have reduced astrocyte cell counts and increased GFAP immunopositive space

Astrocytes in control eyes were longitudinally oriented, more regular, and had a more organized template than myopic eyes, particularly in the peripapillary area (Fig 5A, center). The number of astrocytes was lower in myopic than in control eyes in the superior, inferior, nasal and temporal peripapillary and peripheral regions (Figure 5B and Figure 5D, all p<0.05). Despite the reduced astrocytes cell counts, the spatial coverage of astrocyte processes quantified as the ratio of GFAP+/Sox9 was significantly greater in the peripapillary but not the peripheral retina of myopes compared to controls (Figure 5C, Peripapillary p<0.001; Periphery P=0.21). There was no significant difference in the temporal retina’s GFAP+/Sox 9 ratio (Peripapillary P=0.16, Periphery P=0.36). The astrocytes in the parafoveal region appeared different depending on the layer which they are found (Figure 6A and 6B). The number of astrocytes in the foveal and parafoveal retina was significantly decreased in the myopic RPC vascular layer (Figure 6C, Fovea P<0.001, Parafovea P<0.001) and the myopic superficial vascular layer (Figure 6E, Fovea P<0.001, Parafovea P<0.001). The GFAP+/Sox 9 ratio was greater in the RPC and superficial vascular plexis of myopic foveas and parafoveas (RPC Figure 6D, Fovea P<0.001, Parafovea P<0.001; superficial Figure 6F, Fovea P<0.001, Parafovea P<0.001). Myopic astrocytes had a less complex branching pattern and a reduction in the number of processes.

**Figure 5:**
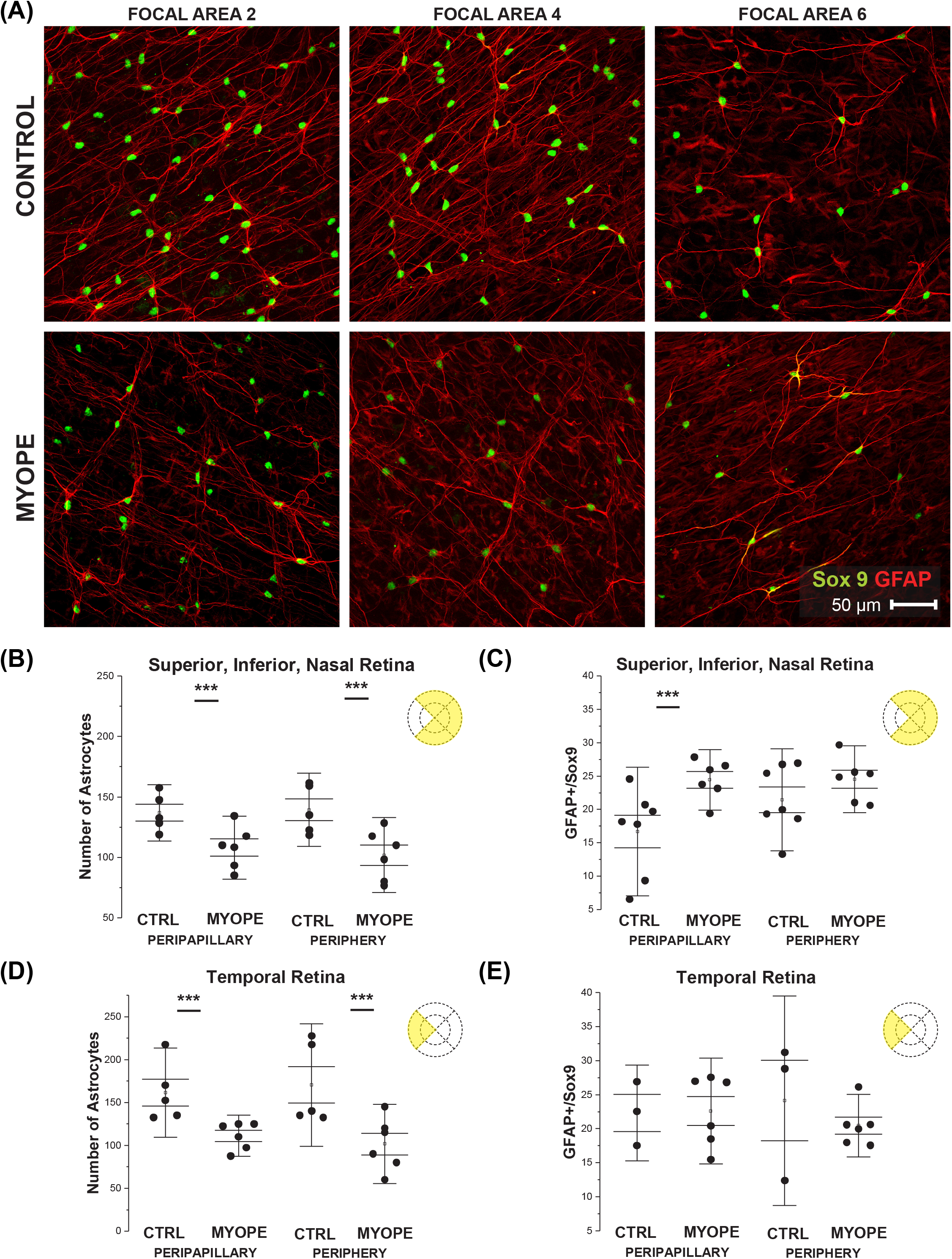
Myopic marmoset retinas show decreased numbers of astrocytes and increased GFAP immunopositive staining in the peripapillary and peripheral retina, excluding the foveal region. **(A)** Representative images of superficial astrocytes in the focal areas 2, 4, and 6 of control (ID tag: C16, age: 268 days, Rx −0.13D) and myopic (ID tag P17, 183 days old, Rx: −7.96D) marmosets. Images are taken at 40x, and astrocytic cell nuclei and bodies/processes in the retinas were labeled with Sox 9 (green) and GFAP (red) markers respectively. **(B)** Shows analyses performed for number of astrocytes in the superior, inferior, and nasal retina. Data shown as a I-graph box plot for control (n=5) and myopic (n=6) marmoset retinas where the box represents SE and whiskers signify SD in (B) through (E). The number of astrocytes decreased significantly in the myopic retina in the superior, inferior, and nasal regions (Peripapillary P<0.001, Peripheral P<0.001) **(C)** The ratio of GFAP immunopositive staining (GFAP+) per Sox-9 labeled astrocyte nuclei (GFAP+/Sox9) increased significantly in the myopic peripapillary region of the superior, inferior, and nasal retina (Peripapillary P<0.001, Peripheral P=0.08). **(D)** The number of astrocytes in the temporal retina decreased significantly in myopic eyes (Peripapillary P<0.001, Peripheral P<0.001), similar to the other regions of the retina. **(E)** The GFAP+/Sox9 ratio did not show significant change between the temporal control and the myopic peripapillary region and periphery region (Peripapillary P=0.16, Peripheral P=0.21).

**Figure 6:**
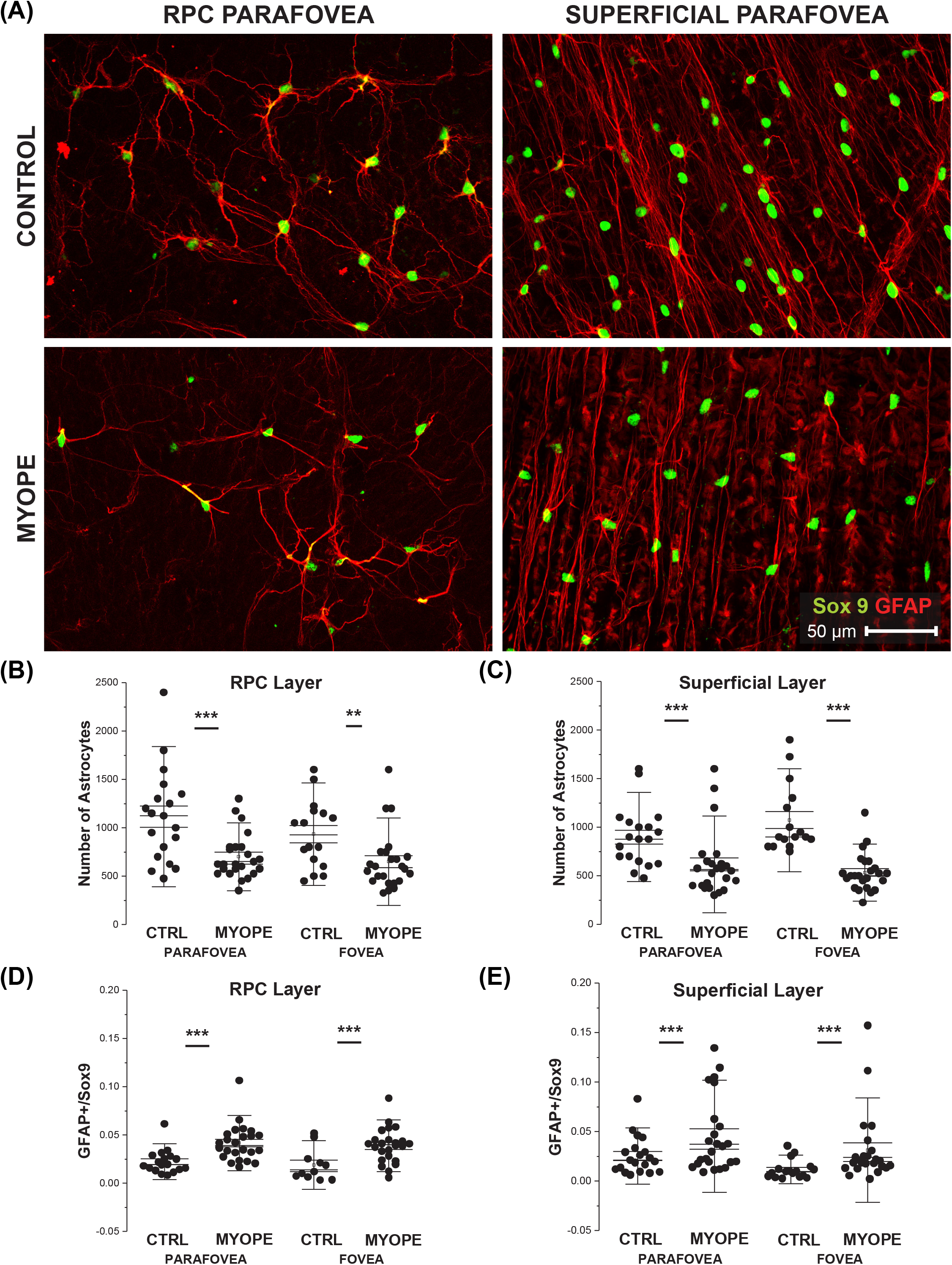
Myopic marmoset retinas showed decreased numbers of astrocytes and increased GFAP immunopositive staining per astrocyte in both the foveal and parafoveal region. **(A)** Representative images of RPC and superficial layer astrocytes in the parafoveal region of control and myopic marmosets (control ID tag: H16 OD, age: 205 days, Rx −0.63D) and myopic (ID tag: P17 OD, age: 183 days, Rx −7.96D). Images were taken at 40x, and astrocyte cell nuclei and body/processes in the retinas were labeled with Sox9 (green) and GFAP (red) markers, respectively. **(B)** Shows analysis performed for the number of astrocytes in the foveal and parafoveal regions (control n=5, myopic n=6). Data is shown as a box plot with SE as the box and SD for whiskers in (B)-(E). The number of astrocytes in the RPC vascular plexus of both the foveal and parafoveal regions decreased significantly in myopic eyes (Fovea P<0.001, Parafovea P<0.01). **(C)** A significant increase in GFAP+/Sox9 ratio was found in the RPC vascular plexus of same areas in the myopic retina (Fovea P<0.001, Parafovea P<0.001). **(D)** The number of astrocytes in the superficial vascular plexus of both the foveal and parafoveal regions decreased significantly in myopic eyes (Fovea P<0.001, Parafovea P<0.001). **(E)** The ratio of GFAP+/Sox 9 was significantly increased in the superficial vascular plexus in myopic foveal and parafoveal retinas (Fovea P<0.001, Parafovea P<0.001).

The extent to which myopic retinal stretch and magnification affected astrocyte cell counts was corrected using a tangential equation. The retinal area for the average myopic retina after correcting for magnification was found to be similar to control eyes (0.64 mm in control eyes, 0.65 mm in myopic eyes), confirming that the effect of image magnification on the cell count calculations was minimal.

### Untreated - but not myopic eyes - exhibit thicker RNFL, less GFAP RFI, and more astrocyte numbers

The averaged values of the parafoveal retinal nerve fiber and ganglion cell layer thicknesses (RNFL and GCL) in control and myopic marmosets are depicted over representative SD-OCT marmoset fundus photographs (Figure 7A, left column). Segmentation calculations are shown in Figure 6A (right column). Representative images of GFAP RFI can be seen in Figure 7B, showing the increase in GFAP+ staining through the thickness of the myopic retina compared to that of the control. The myopic RNFL was significantly thinner in the parafovea compared to controls (Figure 7C: p<0.05). The relative frequency index (RFI) of GFAP was significantly increased in the myopic peripheral superior, inferior, and nasal retinas (Figure 7D: Peripapillary P=0.68, Periphery P<0.05). The GFAP RFI was significantly increased in the myopic peripheral temporal retina (Figure 7E: Peripapillary P=0.10, Periphery P=0.03). GFAP RFI was significantly increased in the myopic parafoveal and foveal retinas (Figure 7F: Parafovea P<0.001, Fovea P<0.05).

**Figure 7:**
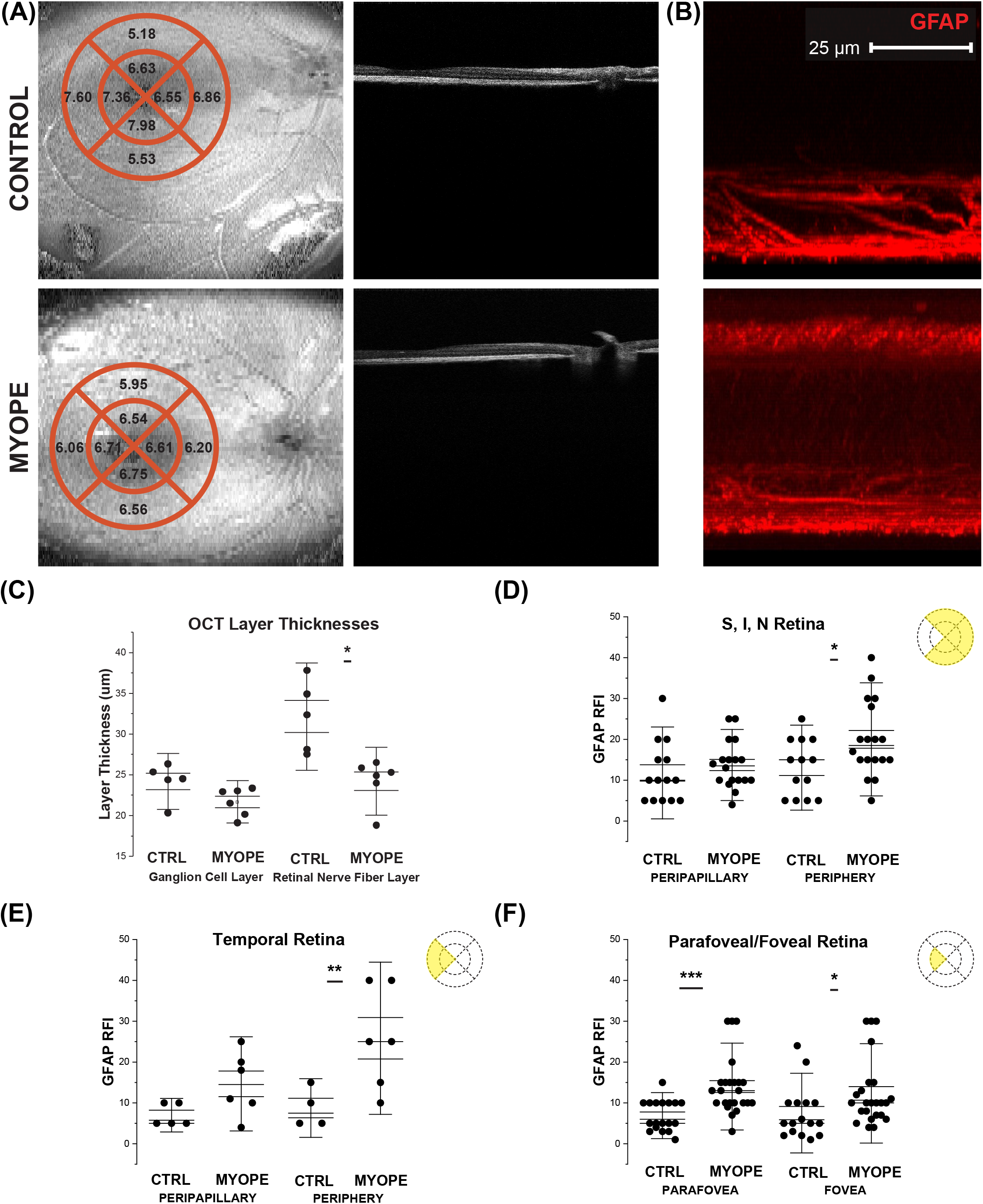
OCTs from the control and myopic marmoset. The graphs show differences in RNFL layer thickness, GCL layer thickness, and correlation between astrocytes and GCL thickness. **(A)** Average ganglion cell layer thicknesses in micrometers (mm) in the quadrants around the fovea of control (top left panel) and myope (bottom left panel) marmosets, as measured with SD-OCT. Representative image of en face optical coherence tomography around the fovea of a control marmoset (top left: ID tag X15, 381 days old, Rx −1.12D) and myopic (bottom left: ID tag O17, 204 days old, Rx −7.28D). Representative cross-sectional scan images of the fovea of the same marmosets can be seen for the control marmoset (top right panel) and myopic marmoset (bottom right panel). **(B)** Representative GFAP RFI images of the superficial retina in a control (top: H16, 205 days old, Rx −1.12D; bottom: P17, 183 days old, Rx −7.96D), showing increased GFAP RFI in the myopic retina compared to the control. **(C)** Analysis of the ganglion cell layer thickness showed that the myopic GCL was no different to the control GCL thickness in the parafoveal retina (P=0.13). However, there is a significant decrease in the myopic RNFL thickness, compared to that of the control RNFL thickness (P=0.04). **(D)** GFAP RFI was significantly increased in the myopic peripheral superior, inferior, and nasal retinas (Peripapillary P=0.68, Periphery P<0.05) **(E)** GFAP RFI was significantly increased in the myopic peripheral temporal retina (Peripapillary P=0.10, Periphery P=0.03). **(F)** GFAP RFI was significantly increased in the myopic parafoveal and foveal retinas (Parafovea P<0.001, Fovea P<0.05).

## DISCUSSION

This study provides evidence of significant changes in three main retinal neurovascular elements and how they relate to each other in a NHP model of lens-induced myopia. Compared to age-matched controls, myopic marmosets lower astrocyte counts in all retinal quadrants, decreased capillary branching in the periphery, increased numbers of string vessels, thinner RNFL, and a lack of association between astrocyte counts and RNFL thickness. In this study, the inner retina’s capillary bed and co-localized astrocytes of marmosets was successfully identified and quantified in the marmoset, a NHP model that has captured the attention of neuroscientists due to its similarity to the human eye. The confocal images obtained from the marmoset retina show a four-layered capillary plexus (radial peripapillary, superficial, intermediate, and deep capillary plexi) and the presence of co-localized astrocytes, similarly to human retinas(Hendrickson et al., 2006, Nickla et al., 2002, Fan et al., 2014).

### Vascular Characterization

The marmoset superficial vascular layer includes main arteries and veins that branch into smaller capillary vessels and form a narrowly stratified capillary network. The quantification analysis described a dense superficial capillary layer that became denser in the deeper layers. The layout and branching pattern identified were comparable to human retinas, indicating a close evolutionary link and confirming marmosets as a good model to study retinal capillary structure and function(Hendrickson et al., 2006). Marmosets with lens-induced myopia had lower capillary branching in the periphery but higher branching in the fovea compared to controls. The myopic decline in peripheral branching suggests a reorganization of the vascular bed as myopic eyes grow and stretch. Reduced capillary branching has been associated with a decreased retinal blood supply in mouse eyes(Bucher et al., 2013, Cheng et al., 2021). There is evidence that as myopic eyes elongate, they can become more prolate in shape with an asymmetric elongation along the horizontal axis(Mutti et al., 2000), stretching the peripheral retina to a greater extent than the central retina. This growth pattern would explain the decreased peripheral but increased foveal branching found in this study. In order to maintain an adequate vascular supply to the increased myopic retinal area, capillary coverage and branching would have to expand. In this study, however, we identified a decrease in peripheral branching. The myopic decrease in peripheral capillary branching may suggest a compromised vascular and metabolic supply to the myopic periphery and potential state of hypoxia(Shih et al., 1993). In fact, there is evidence in the literature that human myopic eyes exhibit non-perfused areas in the far retinal periphery(Hollo et al., 1996). The decrease in capillary branching might also indicate a form of capillary regression, which is known to lead to string vessel formation. String vessels are non-functional capillary strands caused by macrophages engulfing apoptotic endothelial cells(Lang et al., 1994). The existence of these apoptotic endothelial cells has been associated with increasing age, decreased vascular endothelial growth factor (VEGF) concentration(Black et al., 1989), hypoxia(Chavez and LaManna, 2003), and altered blood flow(Rivard et al., 2000, Rivard et al., 1999, Buee et al., 1994, Tilton et al., 1985). In this study, myopic eyes had a greater number of string vessels than controls in the parafoveal retina, suggesting that myopic retinal capillaries are experiencing capillary regression and string vessel formation(Tilton et al., 1981, Tilton et al., 1985, Buee et al., 1994, Kuwabara et al., 1961). While high myopic eyes with thinner choroids have been described to have lower aqueous VEGF concentration(Chen et al., 2015), the relationship between string vessel formation, VEGF and myopia progression remains unexplored.

### Astrocyte Characterization

Studies on astrocyte distribution in the retinal plexi have been done in mice(Cooper et al., 2016, Fernandez-Sanchez et al., 2015), mammals(Bussow, 1980, Cooper et al., 2016, Hollander et al., 1991, Provis et al., 2000, Rungger-Brandle et al., 1993), cats(Hollander et al., 1991) and humans(Schnitzer, 1988, Uga et al., 1974, Karschin et al., 1986), and describe mostly stellate shaped astrocytes in the outermost retinal periphery and more elongated in the central retina(Bussow, 1980, Ogden, 1978, Uga et al., 1974, Liang et al., 2012). The astrocytes observed in marmoset retinas in this study were also stellate in the fovea and periphery, and generally elongated in shape. Astrocytes in the primate retina are proportional in number to the thickness of the RNFL, and have the highest density at the optic nerve head(Rungger-Brandle et al., 1993). This study findings suggest that astrocytes densely populate the marmoset retina, and maintain and establish interactions with endothelial cells in the capillary walls, as previously described in other primates(Hendrickson et al., 2006, Provis et al., 2000). There were similar astrocyte numbers across the control marmoset retina, suggesting that astrocyte growth and development across the retina is fairly symmetrical, similarly to macaque, cat and rabbit retinas(Schnitzer, 1988, Kolb, 1995, Gariano et al., 1994). The length of the astrocyte processes and morphology differed as astrocytes spread out across the retina, suggesting a physiological variation in astrocyte efficiency and function(Rungger-Brandle et al., 1993). Myopic eyes exhibited lower astrocyte numbers than controls in the peripapillary and periphery of myopic eyes, and these numbers remained significant after correcting for magnification. Therefore, ocular magnification secondary to myopic stretch alone cannot explain the decrease in astrocytes identified in this study. This decrease in astrocyte counts suggests a reorganization and redistribution of the astrocyte template as a consequence of myopia development and sustained mechanical stress on astrocyte cell structure. Myopic growth is not symmetric and tends to occur to a greater extent in the periphery. Measurements of peripheral eye length would confirm whether peripheral scaling may be responsible for the lower numbers in the periphery. The lower astrocyte counts might also be related to the known association between astrocyte numbers and RNFL thickness (Lin CR, et al. IOVS 2021;62.8:ARVO E-Abstract), which in this study and others was found to be thinner. There is evidence in the literature of astrocytes increasing GFAP expression during reactive gliosis and remodeling in various non-uniform ways(Sun and Jakobs, 2012). In this study, a significant increase in GFAP immunopositive staining was observed in the myopic parafovea, suggesting a mild astrocyte activation and a potential compromised glial support to the ganglion cells of myopic eyes.

### Astrocytes and Capillaries

The existence and distribution of retinal astrocytes is closely related with the presence and distribution of retinal capillaries(Slavi et al., 2018). Animal models of oxygen-induced retinopathy have described a reduced astrocytic network and highlighted the importance of astrocytes in the formation of retinal revascularization(Bucher et al., 2013, Jiang et al., 1995). The decrease in astrocyte numbers observed in all areas of the myopic marmoset retina, in conjunction with the decrease in capillary branch points, supports the hypothesis that astrocytes might be affected by the vascular changes found. Whether the capillary changes may be leading to hypoxia or decreased perfusion, and how that affects astrocytes remains to be assessed(Bucher et al., 2013). This study did not evaluate retinal oxygenation, but it has been assessed by others: retinal arteriole and arterio-venous oxygen saturation is significantly decreased in highly myopic eyes with and without myopic retinopathy, compared to controls(Zheng et al., 2015). If this was confirmed, astrocytes may be responding to changes in oxygen demand and might be involved in the mechanisms leading to retinal myopic complications.

#### RNFL

Ocular pathologies exhibiting GCL and RNFL thinning with disease progression include glaucoma(Wang et al., 2009), macular degeneration(Lee and Yu, 2015), optic neuritis(Kupersmith et al., 2016), and Alzheimer’s disease(Lopez-de-Eguileta et al., 2020). In this study, the RNFL was significantly thinner in myopic marmoset eyes and remained significant after correcting for the effects of magnification secondary to myopic stretch, suggesting that myopia affects ganglion cell axon distribution, which has been described by others in human eyes(Grytz et al., 2020, Lee and Yu, 2015, Seo et al., 2017, Chen et al., 2020, Jonas et al., 2020).

#### PUTTING EVERYTHING TOGETHER

The results from this study suggest that myopic eye growth affects the architectural template of three key neurovascular elements: capillaries, astrocytes and ganglion cells. The restructuring and reorganization observed in NHP myopic eyes may reflect a dynamic adaptation to the changing environment triggered by myopic eye growth and stretch, and be part of a beneficial adaptive chronic response to support neurons during ocular growth(Maki et al., 2013). Alternatively, it could represent a detrimental response indicating the beginning of a compromised structural vascular support to the inner retina affecting vascular and astrocyte function leading to an alteration in the ability to regulate local ions, neurotransmitters and metabolites, and affect neural function(Maki et al., 2013). The changes in astrocyte morphology and numbers found in the myopic retina suggest that myopic retinal stretch triggers an adaptive response on astrocytes, which might be lengthening their cell bodies and axons in an effort to provide adequate structural and functional support to the vasculature and RGCs. This type of response has been described before in astrocytes exposed to artificially induced mechanical tension, which formed living scaffolds to guide neuroregeneration (Katiyar et al., 2017). However, since astrocytes and retinal capillaries are crucial to maintain RGC axonal viability(Gould et al., 1998, Okano et al., 2012), the reduced astrocyte numbers observed in myopic eyes, along with the increased GFAP immunopositive staining, reduced capillary branching and increased number of string vessels might be affecting neural function. In particular, the increased GFAP immunopositive staining identified in the peripapillary and parafoveal myopic retina, along the RNFL thinning observed in myopic eyes might be a sign of early glial reactivity. The space occupied by neural axons lost due to injury or degeneration is generally filled by a glial scar, and predominantly involves hypertrophic astrocyte processes(Cooper et al., 2016). Astrocytes can become reactive to mechanical and chemical stimuli like elevated intraocular pressure, increased mechanical tension(Katiyar et al., 2017), and excitotoxicity(Sun and Jakobs, 2012). These changes reflect the biphasic nature attributed to astrocytes(Maki et al., 2013), which includes degenerative and regenerative properties at different phases after retinal stress or injury.

In conclusion, this study confirms the feasibility of the marmoset as an experimental model to study the retinal neurovascular unit. The aim was to evaluate in detail the effect of myopic eye growth on the structure and distribution of three key neurovascular elements: capillaries, astrocytes and ganglion cell/retinal nerve fiber layer thickness. This study confirms that myopic eyes without pathology exhibit changes in all three neurovascular elements, suggesting that the neurovascular unit may be affected by myopic mechanical stretch and elongation. Whether this neurovascular adaptation is beneficial or harmful, and whether its function is altered, remains to be investigated.

## METHODS

### Marmoset model of myopia

Eleven juvenile marmoset eyes were studied, six of which were induced with myopia by imposing hyperopic defocus using full field negative single-vision soft contact lenses (−5D). For 80% statistical power to detect effect size, a sample size of n=8, or n=4 in each group, was required. All animal care, treatment and experimental protocols were approved by the SUNY College of Optometry Institutional Animal Care and Use Committee (IACUC), the ARVO statement for the use of animals in ophthalmic and vision research, the US National Research Council’s Guide for the Care and Use of Laboratory Animals, the US Public Health Service’s Policy on Humane Care and Use of Laboratory Animals, and Guide for the Care and Use of Laboratory animals. Identification, age, axial length and refractive error of control and myopic marmosets are listed in Table 1.

**Table 1.**
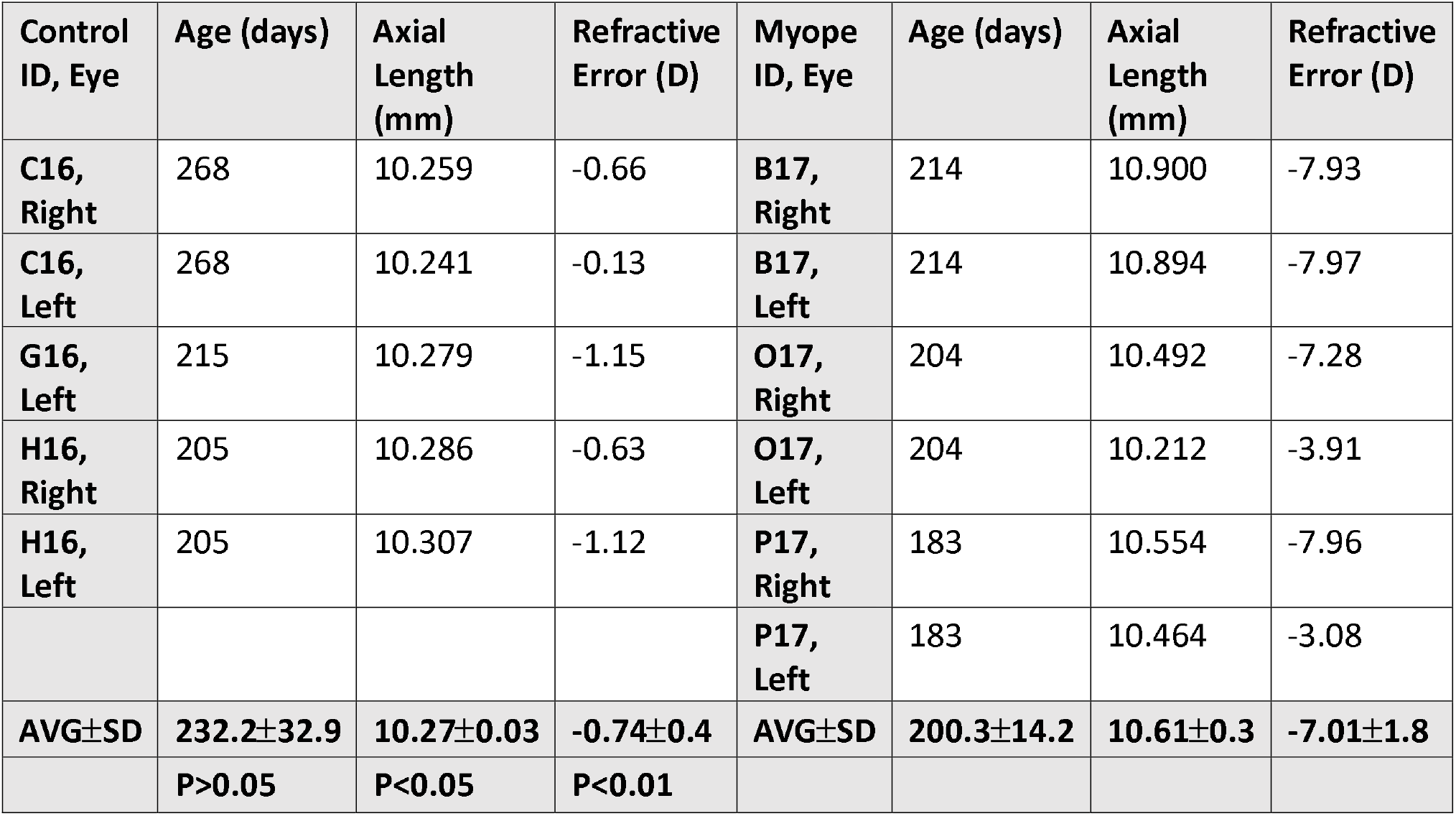
Marmoset identification with age (days) at time of enucleation (ID). In each of the six marmosets, both right (R) and left (L) eyes were enucleated for a total of 11 eyes. The significance between control and myopic group age was not significant (p>0.05); significantly different between axial length (p<0.05); and significantly different between refractive error (p<0.01). Only 11 of the eyes were suitable for immunohistochemical analysis. All useable retinal sections were mounted for confocal microscopy.

| Control ID, Eye | Age (days) | Axial Length (mm) | Refractive Error (D) | Myope ID, Eye | Age (days) | Axial Length (mm) | Refractive Error (D) |
| --- | --- | --- | --- | --- | --- | --- | --- |
| C16, Right | 268 | 10.259 | -0.66 | B17, Right | 214 | 10.900 | -7.93 |
| C16, Left | 268 | 10.241 | -0.13 | B17, Left | 214 | 10.894 | -7.97 |
| G16, Left | 215 | 10.279 | -1.15 | O17, Right | 204 | 10.492 | -7.28 |
| H16, Right | 205 | 10.286 | -0.63 | O17, Left | 204 | 10.212 | -3.91 |
| H16, Left | 205 | 10.307 | -1.12 | P17, Right | 183 | 10.554 | -7.96 |
|  |  |  |  | P17, Left | 183 | 10.464 | -3.08 |
| AVG±SD | 232.2±32.9 | 10.27±0.03 | -0.74±0.4 | AVG±SD | 200.3±14.2 | 10.61±0.3 | -7.01±1.8 |
|  | P>0.05 | P<0.05 | P<0.01 |  |  |  |  |

Treatment started at 10 weeks of age (72.0±5.0 days) following our established protocol(Benavente-Perez et al., 2012, Benavente-Perez et al., 2014, Benavente-Perez et al., 2019). Lenses were inserted daily in the morning between 8-10am when lights were turned on in the animal room (700 lux) and removed 9hrs later at lights off each day (9 hours light/15 hours dark)(Benavente-Perez et al., 2014, Troilo and Judge, 1993). Contact lenses had 6.5 mm diameter, 3.6/3.8 mm base curve, were made of methafilcon A (55% water content, DK: 17), fit 0.10 mm flatter than the flattest keratometry measurement, and assessed using an ophthalmoscope. No corneal complications were observed in any of the animals treated in this or earlier studies with marmosets(Benavente-Perez et al., 2014, Troilo and Judge, 1993).

Cycloplegic refractive error (Rx, Nidek ARK-700A autorefractor, Nidek Co., LTD, Aichi, Japan), ocular axial length (AL, Panametrics, NDT Ltd, Waltham MA) and spectral domain optical coherence tomography (SD-OCT, Bioptigen SD-OCT; 12×12 mm^2^, 700 A-scans/B-scan x 70 B-Scans x 5 Frames/B-scan) were performed at baseline and end of treatment prior to enucleation. RNFL and GCL thickness were measured using the SD-OCT under anesthesia (alphaxalone, 15mg/kg, IM), and segmented and quantified using The Iowa Reference Algorithms v3.6 (Iowa Institute for Biomedical Imaging).

### Enucleation, dissection, and flat mount preparation

At the end of treatment, eyes were enucleated and placed in phosphate buffer saline (PBS) (ThermoFisher). Dissected retinas were fixed in Para-Formaldehyde (PFA) 4% in PBS (Santa Cruz Biotechnology) for 40 minutes, washed five times for 30 minutes each with PBS and incubated with 5% normal goat serum (ThermoFisher) and 0.5% TritonX (Sigma Aldrich) blocking buffer to avoid non-specific antibody binding. Following blocking, the retina was incubated with primary antibodies diluted in blocking buffer at 4°C for 3 days. The primary antibodies used in this study were isolectin-Alexa 488 (1:100) (ThermoFisher), mouse anti-GFAP (1:500) (Sigma Aldrich), and rabbit anti-Sox 9 (1:1000) (Sigma Aldrich). After the incubation period, the retinas were washed five times for 30 minutes each with PBS and incubated with goat-anti mouse secondary antibody conjugated with Texas Red (1:500) (ThermoFisher) and goat-anti rabbit 647 (1:500) (ThermoFisher). The SuperFrost slides (ThermoFisher) were cleaned with ethanol. Retinas were inspected for any signs of debris, and consistent tissue thickness achieved by pinching and cutting vitreal remains. Retinas were plated and cover slips were placed on objectives with DAPI-Mounting medium (Vector Laboratories), permitted to self-seal and stored at −20°C.

### Confocal Microscopy and Image Acquisition

The immunochemical samples were imaged using the Olympus FV1200 MPE confocal microscope. Sixteen images (640 μm x 640 μm along the horizontal plane, and 10 μm along the vertical plane) were taken from each of the eleven retinas imaged. Multi-plane z-series were collected using a 20x objective, with each section spaced 1 μm apart. These 10 sections were processed by the confocal microscope to form a single z-stack of images subtending the whole specimen. Images were processed using Fiji software.

The structure and distribution of astrocytes and co-localized blood vessels were assessed by imaging all four retinal quadrants (temporal, nasal, superior, and inferior) in the periphery, peripapillary, and parafoveal retina. This regional analysis was performed with the goal to identify local changes that might occur in myopic eyes due to their assymetric eye growth pattern. A representation of the locations evaluated can be seen in Figure 1A, which is a composite of 60 individual frames of a control marmoset retina (ID: X15, age: 381 days, refractive error: −1.12D) taken at 4x magnification visualized with Isolectin-488 using the Olympus confocal microscope. To ensure a detailed analysis, images were acquired at higher magnification in focal area 3 for the peripapillary region, and focal area 6 for the peripheral region. An independent foveal analysis was performed on the foveal and parafovea regions (boxes 1 and 2, Figure 1B). Detailed images of the foveal (Figure 1C) and parafoveal (Figure 1D) regions show the difference in anatomy between the two areas. Images were also acquired at focal areas 1-6 starting from the optic nerve head, as shown in Figure 1A, which represent focal distances away from the optic nerve head.

### Image Analysis

#### Blood vessel analysis

Blood vessel branching points, as well as the presence and number of string vessels per image frame were manually counted for each frame on branches of all orders and converted to number of branch points/mm^2^ and number of string vessels/mm^2^, respectively. The branch points and string vessels were quantified in the superficial capillary plexus.

#### Astrocyte quantification

The number of astrocyte nuclei was counted in every image frame using Fiji cell counter function and converted to astrocytes/mm^2^. The image was split into its color channels to identify the red channel corresponding to GFAP. Fiji automatically analyzed GFAP immunopositive staining density, which was converted to a ratio of GFAP+/Sox 9 for our analysis. Astrocyte spatial coverage was quantified as the ratio of GFAP immunopositive staining (GFAP+) per Sox 9 positive astrocytes (GFAP+/Sox9) in the superficial plexus to represent the amount of GFAP+astrocytic processes per astrocyte nuclei.

The increased GFAP+/Sox 9 ratio in the myopic retinas in our flatmount images probed us to perform three-dimensional reconstructions of the retinal tissue to see how GFAP distributes through the marmoset retina’s depth. The “3D stack function” on Fiji constructed a three-dimensional image from the flatmounts by flipping the z-stacks 90° on the X axis. We drew a vertical line horizontally across these 3D reconstruction images, of the same length and in the same area for all eyes. The “plot profile” function on Fiji then created a two-dimensional graph of pixel intensities along the line, with the x-axis representing distance in μm and the y-axis representing gray scale intensity. The “relative frequency index” (RFI was then calculated by averaging the values of minimum GFAP intensity subtracted from the average maximum GFAP intensity in all areas. This value was gathered in the myopic and control peripapillary and peripheral retina, where the most myopic GFAP immunopositive staining was found.

#### Correction for Magnification

The extent to which myopic retinal stretch affected the capillary and astrocyte calculations was determined using a tangential equation. For the OCT calculations, the diameter of the ETDRS grid circle was adjusted for ocular magnification using the formula: *R*_N_= (*R* ×*AL*)/9.62, where R is the radius of the ETDRS grid and AL is the axial length of the marmoset. The value 9.62 is the average axial length in millimeters of adult control marmosets in our lab.

### Statistical Analysis

Data was assessed for normality and analyzed using an analysis of variance (ANOVA) at the level of α= 0.05 using OriginPro 2021b software (OriginLab, Northampton, Massachusetts, USA).

Associations between the number of astrocytes and ganglion cell layer thickness was studied using multiple regression.

## Acknowledgments

a. Stefanie Wohl, Harrison Feng, Rita Nieu, Amy Pope, Victor Lin, Ana Nour.

## Data Availability

The data that support the findings of this study are available from the corresponding author upon reasonable request.

